# Defining levels of dengue virus serotype-specific neutralizing antibodies induced by a live attenuated tetravalent dengue vaccine (TAK-003)

**DOI:** 10.1101/2020.09.23.308817

**Authors:** Laura J. White, Ellen Young, Mark Stoops, Sandra Henein, Ralph S. Baric, Aravinda M. de Silva

**Author notes:** Department of Marine Sciences, University of North Carolina at Chapel Hill (Mark Stoops). Corresponding author: Laura White (LW).

## Abstract

The four dengue virus serotypes (DENV1-4) infect several hundred million people each year living in tropical and sub-tropical regions. Clinical development of DENV vaccines is difficult because immunity to a single serotype increases risk of severe disease during a second infection with a new serotype. Leading vaccines are based on tetravalent formulations to induce simultaneous and balanced protective immunity to all 4 serotypes. TAK-003 is a tetravalent live attenuated dengue vaccine candidate developed by Takeda Vaccines Inc, which is currently being evaluated in phase 3 efficacy trials. Here, we use antibody depletion methods and chimeric, epitope transplant DENVs to characterize the specificity of neutralizing antibodies in dengue-naïve adults and non-human primates immunized with TAK-003. Our results demonstrate that TAK-003 induced high levels of DENV2 neutralizing antibodies that recognized unique (type-specific) epitopes on DENV2. In contrast, most vaccinated subjects developed lower levels of DENV1, DENV3 and DENV4 neutralizing antibodies that mainly targeted epitopes that were conserved (cross-reactive) between serotypes. We conclude that the DENV2 component in the vaccine is immunodominant because of the high levels of serum neutralizing antibodies targeting type-specific epitopes. We also conclude that DENV1, 3 and 4 vaccine components are less immunogenic because most study subjects did not develop type-specific serum neutralizing antibodies to these serotypes. While DENV vaccine development has been guided by the presence of neutralizing antibodies to each serotype as a benchmark, our results indicate that the presence of neutralizing antibodies alone are not a reliable indicator of the immunogenicity of each vaccine component.

**Author summary:** The development of tetravalent dengue vaccines has been guided by neutralizing antibodies to each serotype as a correlate of safe and effective vaccine induced immunity. However, the absolute levels of neutralizing antibodies to each serotype has proven to be an unreliable correlate of protection. Levels of antibodies to epitopes that are unique to each serotype, which are measures of immunity independently stimulated by each vaccine component, rather than total quantity of neutralizing antibodies, are likely to be better correlates of protection. Here, we mapped the specificity of antibodies induced by the Takeda tetravalent dengue vaccine TAK-003 in monkeys and humans with no prior immunity to dengue. The TAK-003 vaccine induces high levels of serotype 2 specific neutralizing antibodies that map to known protective epitopes. In contrast, the serotype 1, 3 and 4 neutralizing antibody responses are lower and mainly consist of cross-reactive antibodies binding to epitopes conserved between serotypes. These heterotypic antibodies, which are most likely derived from the serotype 2 component, may not provide long term protection in vivo.

## Introduction

The four dengue virus serotypes (DENV1-4) infect several hundred million people each year living in tropical and sub-tropical regions (1-3). Clinically, DENV infections can be inapparent or present as a febrile illness that may or may not progress to severe dengue hemorrhagic fever and shock syndrome. Primary infection with a DENV serotype leads to long-term protective immunity against the homologous serotype and transient cross protection against other serotypes (4, 5). DENV serotype-specific neutralizing antibodies, which circulate for decades if not longer, are correlated with durable protection against re-infection by the same serotype (6, 7). Secondary DENV infections with heterologous serotypes pose increased risk of progression to severe dengue hemorrhagic fever and shock syndrome. The ability of some DENV cross-reactive antibodies to promote the entry of heterologous serotypes into Fc receptor-bearing target cells is widely supported as the initiating event that culminates in severe disease (8, 9). To minimize the risk of DENV vaccines inducing antibodies that enhance DENV infections, leading vaccines are based on tetravalent formulations to induce simultaneous and balanced protective immunity to all 4 serotypes. However, in practice, it has been challenging to achieve balanced replication and immunity with tetravalent DENV vaccines. Here we characterize the properties of neutralizing antibodies induced by a leading live attenuated tetravalent DENV vaccine (TAK-003) to determine the contribution of each vaccine component to the serotype-specific neutralizing antibody response.

After a primary DENV infection the durable protective immune response is directed to serotype-specific epitopes on the infecting virus, leaving individuals fully susceptible to a second infection with a new serotype. After a second infection with a new serotype, the protective immune response is mainly directed to epitopes conserved between serotypes, and clinically significant tertiary DENV infections are rare (10, 11). While the immune mechanism responsible for serotype cross-protective immunity after a second infection have not been full elucidated, immunological memory from the first infection is critical for establishing durable cross-protective immunity. Recent studies indicate that reactivation of memory cells by a second infection selects for high affinity, cross-reactive antibodies and T cells responsible for cross-protective immunity (12-14). These studies are relevant to understanding how live attenuated tetravalent DENV vaccines will protect people (15). In people who have experienced a DENV infection before receiving the vaccine, even a poorly balanced vaccine dominated by one vaccine serotype is likely to induce cross protective immunity by activating memory. In DENV naïve individuals with no immune memmory, the protective immune response to each serotype will strongly depend on how well each vaccine components replicates.

Several DENV vaccines are under development, including two live attenuated tetravalent DENV vaccines (TAK-003 developed by Takeda Vaccines Inc and TV-003 developed by the US National Institutes of Health) currently in phase III clinical trials and one live attenuated tetravalent vaccine, Dengvaxia, developed by Sanofi Pasture that has been licensed for use in children with pre-existing immunity to DENV (16-18). In children with no immunity to DENVs, Dengvaxia was poorly efficacious and the vaccine increased the risk of severe dengue disease upon exposure to wild type DENV infections. The poor performance of Dengvaxia in dengue-naïve children is mainly due to unbalanced replication of vaccine components, where the DENV4 component is replication- and immune-dominant compared to the other three serotypes (19-23).

TAK-003 developed by Takeda is based on an attenuated DENV2 strain and 3 additional chimeras of the attenuated DENV2 strain expressing the envelope proteins of DENV1, 3 and 4. This candidate, which has been shown to be safe and immunogenic in phase 1 and 2 trials, is currently being evaluated in a large multi-country phase 3 efficacy trial (24, 25). In preclinical and early clinical studies, the DENV2 component of the vaccine was observed to replicate better than the other three chimeric components (26, 27) (28). In people with no pre-existing immunity DENVs, TAK-003 stimulated higher levels of neutralizing antibody to DENV2 compared to the other three serotypes (24, 25).

While TAK-003 stimulates lower levels of neutralizing antibodies to DENV1, 3 and 4, it is unclear if these antibodies target epitopes that are conserved across DENV serotypes or epitopes that are unique (type-specific) to each serotype. Neutralizing antibodies directed to type-specific epitopes are indicative of a vaccine component replicating and independently stimulating antibodies to a DENV serotype. Here, we describe the levels of DENV serotype-specific and cross-reactive neutralizing antibodies in dengue-naïve adults and non-human primates immunized with TAK-003.

## Methods

### Cell lines and viruses

C6/36 *Aedes albopictus* mosquito cells were used to grow DENV1-4 to use in ELISA and virus neutralization assays. Cells were maintained in minimal essential medium (MEM; Gibco) at 32°C. Vero-81 mammalian cells (American Type Culture Collection; CCL-81) were used to grow DENV1-4 for the generation of purified antigens. Vero cells were maintained in Dulbecco’s modified Eagle’s medium-F12 (DMEM-F12) at 37°C. All growth and maintenance media used were supplemented with 5% fetal bovine serum (FBS), 100 U/mL penicillin, 100 mg/mL streptomycin, 0.1 mM non-essential amino acids (Gibco), and 2 mM glutamine. All cells were incubated in the presence of 5% CO2. The 5% FBS was reduced to 2% to make infection medium for each cell line. DENV1 (American genotype; strain West Pac74), DENV2 (Asian genotype; strain S-16803), DENV3 (Asian genotype; strain CH-53489), and DENV4 (American genotype; strain TVP-376) (provided by Robert Putnak, Walter Reed Army Institute of Research, Silver Spring, MD) were used for both binding enzyme-linked immunosorbent assays (ELISAs), neutralization assays, and depletion assays, unless otherwise noted. DENV1-4 coupled to Dynabeads in depletion assays were grown in Vero-81 cells and purified by tangential flow concentration and density gradient ultracentrifugation. All the recombinant chimeric viruses and WT infectious clones used here were constructed at UNC using a cuadripartite complementary DNA clone, as reported earlier (29).

### Ethical Statement

This study used deidentified human samples provided by Takeda Vaccines Inc. The UNC IRB classified the research as exempt from local IRB review. Serum samples were collected by Takeda Vaccines Inc. in the completed Phase 2 clinical trial DEN-205 (ClinicalTrials.gov Identifier: NCT02425098). DEN-205 was conducted in accordance with Institutional Review Board regulations in Singapore, the Declaration of Helsinki, International Council for Harmonization of Technical Requirements for Registration of Pharmaceuticals for Human Use and Good Clinical Practice (ICH-GCP) guidelines, and applicable regulatory requirements.

### Source of non-human primates serum samples

The sera used in this study was received from Takeda Vaccines Inc. We analyzed day 180 serum samples from 30 adult DENV sero-negative Indian rhesus macaques that had been immunized with the tetravalent TAK-003 vaccine, or infected with wild type DENV1, DENV2 or DENV3. (S1 Table).

### Source of human serum samples

Subjects in the Phase 2 clinical trial DEN-205 were randomly assigned 1:1 to receive a single dose of either TDV or HD-TDV on Day 1 of the trial. Random assignment was stratified based on dengue IgG status (positive or negative) at screening, with a maximum of 200 subjects to be enrolled per dengue IgG status group (28). We received from Takeda 30 baseline seronegative serum samples, including 14 TDV and 16 HD-TDV. Here we analyzed sera (N=30) from seronegative subjects collected 180 days after vaccination, and sera (N=16) from seropositive subjects collected on day 0 before vaccination (S2 Table). The study reported here is not part of the primary or secondary objectives of the trial.

### Vaccine formulations

The DEN-205 subjects included in this study received one of two vaccine formulations. The HD-TDV formulation contained 2 × 104 plaque-forming units (pfu) of TDV-1, 5 × 104 pfu of TDV-2, 1 × 105 pfu of TDV-3, and 3 × 105 pfu of TDV-4. The TDV formulation contained identical dosages of TDV-1, TDV-3, and TDV-4, but a lower dosage of TDV-2 (5 × 103 pfu) (S3 Table) (28). The experimental vaccine received by NHPs consisted of 3.17×10^4^ plaque-forming units (pfu) TDV-1, 9.72 x10^4^pfu TDV-2, 3.75 x10^5^ pfu TDV-3 and 3.65 x10^5^ pfu TDV-4.

### Antibody depletions

We have developed antibody depletion methods to separate antibodies in polyclonal sera into cross-reactive (CR), which are directed to epitopes on the virion that are conserved among dengue virus serotypes, and serotype-specific Abs (TS), which bind unique epitopes on each serotype (11, 23, 30). In principle, Abs with specific binding properties are removed from sera by incubation with purified whole virion antigen of the desired serotype covalently linked to polystyrene or magnetic beads. The properties of Abs in undepleted and Ab-depleted serum are then assessed by ELISA and mFRNT assays.

Samples from NHP were depleted using purified DENV or BSA control protein adsorbed onto Polybeads polystyrene microspheres (Polysciences, Inc.) as reported earlier (11, 23). Samples from vaccinated human subjects were depleted following a modified depletion protocol using magnetic beads and magnetc separation. Briefly, Magnetic Dynabeads M280 Tosyl activated (DB) (Invitrogen 14203, 14204) were first covalently linked to the dengue cross-reactive monoclonal antibody 1M7 following manufacturer’s protocol (50 ug 1M7 per 5 mg DB). DB-1M7 complex was blocked with BSA in PBS and then washed with 0.1 M 2-(*N*-morpholino) ethane sulfonic acid (MES) buffer, and then incubated for 1 h at 37°C in separate with purified DENV1, DENV2, DENV3, DENV4 or Bovine Serum Albumin (BSA) antigen control at a ratio of 100 ug of antigen per 5 mg DB. After 3 PBS washes the bound antigen was cross-linked to the DB-1M7 complex with 2% PFA for 20 min at room temperature. Serum samples were diluted 1:10 and incubated with the Antigen-DB complex for 1 h at 37°C with end-over-end mixing under 3 conditions: a) BSA-DB as undepleted control (total Nabs, TS+CR) (UND), b) Heterologous serotypes-DB to remove CR Abs and leaves TS NAbs, (HET), and c) Homologous serotype-DB, as indicator of depletion efficiency (HOM). Abs bound to beads were magnetically separated from the depleted serum. For the analysis of clinical samples, a paired depletion approach was used, where after DV1-DB+DV3-BD heterologous depletions (HET) of serum, the remaining titers to DV2 and DV4 represented the fraction of TS NAbs to DV2 and DV4 respectively. On the other hand, remaining Neut_50_ titers to DV1 and DV3 after depletions with DV2-DB+DV4-DB (HET) represent the fraction of TS NAbs to DV1 and DV3. Homotypic depletion (HOM) efficiency was determined for each serotype by ELISA and mFRNT. For the analysis of preclinical samples, HET depletion was performed with DENV2 for measuring TS NAbs to DENV1 and DENV3. To measure TS NAbs to DENV2, HET depletion was performed with a mix of DB conjugated to DENV1, DENV3 and DENV4. Three rounds of depletion were usually needed for successful removal of >80% of homologous dengue Abs.

### Analysis of TS NAbs

Neut_50_ titers in the undepleted, homotypic-depleted and heterotypic-depleted sera were determined by mFRNT. The fraction of Abs in the sera that were TS was calculated using two independent methods. Method 1 used the formula: %TS NAbs = [Neut_50_ HET depletion – Neut_50_ HOM depletion]/[Neut_50_ BSA depletion – Neut_50_ HOM depletion] × 100, as reported earlier (23). Method 2 used log10 transformed values of each Neut_50_ titer to calculate levels of TS NAbs using the equation = [(log10 Neut_50_ HET depletion/log10 Neut_50_ BSA depletion) x100]. A value above 55 was selected as cutoff for evidence of TS NAbs. Other criteria for inclusion of samples in TS analysis using Method 2 were heterologous depletion efficiency >90% and samples with BSA-depleted Neut_50_ titer of <40 were consider TS negative.

### ELISA

Antigen capture ELISA was used to confirm efficient depletion of targeted antibodies. Briefly, purified antigen was plated at 100ng/well in a 96 well ELISA plates overnight at 4°C. Plates were blocked with 3% (vol/vol) normal goat serum (Gibco, Thermo Fisher, USA)-Tris buffered saline (TBS) −0.05% (vol/vol) Tween 20 (blocking buffer). Depleted sera were diluted and added to the antigen coated ELISA plates. Alkaline phosphatase-conjugated secondary Abs were used to detect binding of sera with p-nitrophenyl phosphate substrate, and reaction color changes were quantified by spectrophotometry at 405nm.

### DENV neutralization assay

To measure neutralizing antibody titers in un-depleted and depleted sera, we used a micro focus reduction neutralization test adapted to a 96-well plate format (mFRNT) as described earlier (31). Briefly, 96-well plates were plated with 2 × 10^4^ Vero-81 cells/well and incubated at 37°C for 24 hours. Serial 3-fold dilutions of each serum were mixed with 50-100 FFU of virus in DMEM with 2% FBS. The virus-Ab mixtures were incubated for 1 h at 37°C and then transferred to the confluent monolayer of Vero 81 cells on the 96-well plates. Following an additional 1 h incubation at 37°C, the monolayers were overlaid with Opti-MEM (Gibco, Grand Island, NY) containing 2% FBS and 1% (wt/vol) carboxymethyl cellulose (Sigma, St. Louis, MO). EC50 values were calculated by graphing % neutralization vs. serum dilution and fitting a sigmoidal dose response (variable slope) using Prism 8 (GraphPad Software, San Diego, CA, USA). Neut_50_ titers represent the dilution at which the serum neutralizes 50% of the infection. Log transformed data from Neu50 values were used to calculate GMT +/-95%CI. Criteria to accept values to be reported were an R^2^ >0.75 and a Hill Slope >0.5 absolute value.

### Titration of wild-type (WT) and chimeric viruses on Vero cells

For titrations of WT and chimeric viruses, viral stocks were diluted 10-fold serially in dilution medium (OptiMEM, Grand Island, NY) supplemented with 2% heat-inactivated fetal bovine serum (HI-FBS; Hyclone Defined) and 1x antibiotic/antimycotic. Following a 1hr infection, cells and inoculum were overlaid with overlay medium: OptiMem (Gibco, Grand Island, NY) containing 5% carboxymethylcellulose, 2% HI-FBS (Hyclone defined), and 1x antibiotic/antibiotic (Gibco, Grand Island, NY). Following a 2 day incubation (to achieve countable foci of infection), cell monolayers were washed with 1xPBS followed by fixation/permeabilization with 4% paraformaldehyde/ 0.01% saponin (v:v). Fixed cell monolayers were blocked in a solution of PBS containing 5% non-fat milk and incubated with a primary antibody cocktail of 4G2 and 2H2 murine mAbs. Following thorough PBS washes, infectious foci were visualized using HRP-goat-anti-mouse secondary antibody (KPL, Gaithersburg, MD), followed by TrueBlue substrate and foci were enumerated and used to calculate infectious titer. Number and size of foci were analyzed with a CTL Immunospot instrument.

### Statistical analysis

Variation between groups was measured by a non-parametric Friedman test with Dunn’s multiple comparisons. P < 0.05 were considered statistically significant.

## RESULTS

### Levels of DENV serotype specific NAbs in nonhuman primates after TAK-003 vaccination

Non-human primates (NHP) have been useful models to evaluate the immunogenicity and protective efficacy of dengue vaccines (26, 32). Here we used NHPs sera from Takeda study DNHP-007 (S1 Table) to characterize antibodies induced by TAK-003 vaccination and WT DENV infection. The monkeys were vaccinated with one (N=4) or two doses (N=4) of the TAK-003 or infected with WT DENV1, 2 or 3 (N=2 per serotype). Serum was collected 180 days after the last vaccine dose or WT DENV infection. The immune sera were pre-incubated with magnetic beads coated with different DENV serotypes to remove DENV serotype cross reactive antibodies, while retaining any type-specific antibodies to the serotype of interest.

As depicted in Fig 1, undepleted sera reflect total levels of NAbs to each serotype (Fig 1A), and HET depleted sera reflect levels of serotype specific NAbs (Fig 1B, HET Depleted). The vaccine induced total NAbs titers >20 to DENV1 in 7/8 animals (GMT= 233), to DENV2 in 8/8 (GMT=4,489), and to DENV3 in 8/8 animals (GMT=145). The titers to DENV2 were significantly higher than those to DENV1 and DENV3. After a second dose of TDV (half-filled symbols), a modest (<4-fold) increases in NAb titers to each serotype was seen in some animals, but there was no significant difference between one and two doses. After WT DENV1, DENV2 or DENV3 infections, mean titers were 10,375, 9135 and 1742 respectively. Fig 1B shows the corresponding Neut_50_ titers after heterologous depletion, representing the fraction of induced NAbs binding to type-specific epitopes on each serotype. All animals showed evidence of DENV2 TS NAbs after HET depletion (GMT=3,077), with the TS fraction for each animal ranging between 39 and 100% of total NAbs. For DENV1, only 1/8 animals had TS NAbs after HET depletion (Neut_50_ = 23). For DENV3, 4/8 had low levels of TS NAbs (GMT=34). Animals infected with WT DENV1, DENV2 and DENV3 developed high levels of NAbs, and HET depletion resulted in minimal change in titers (Fig 1B), indicating that a large fraction of NAbs induced by a WT DENV infection in NHP are TS.

**Figure 1.**
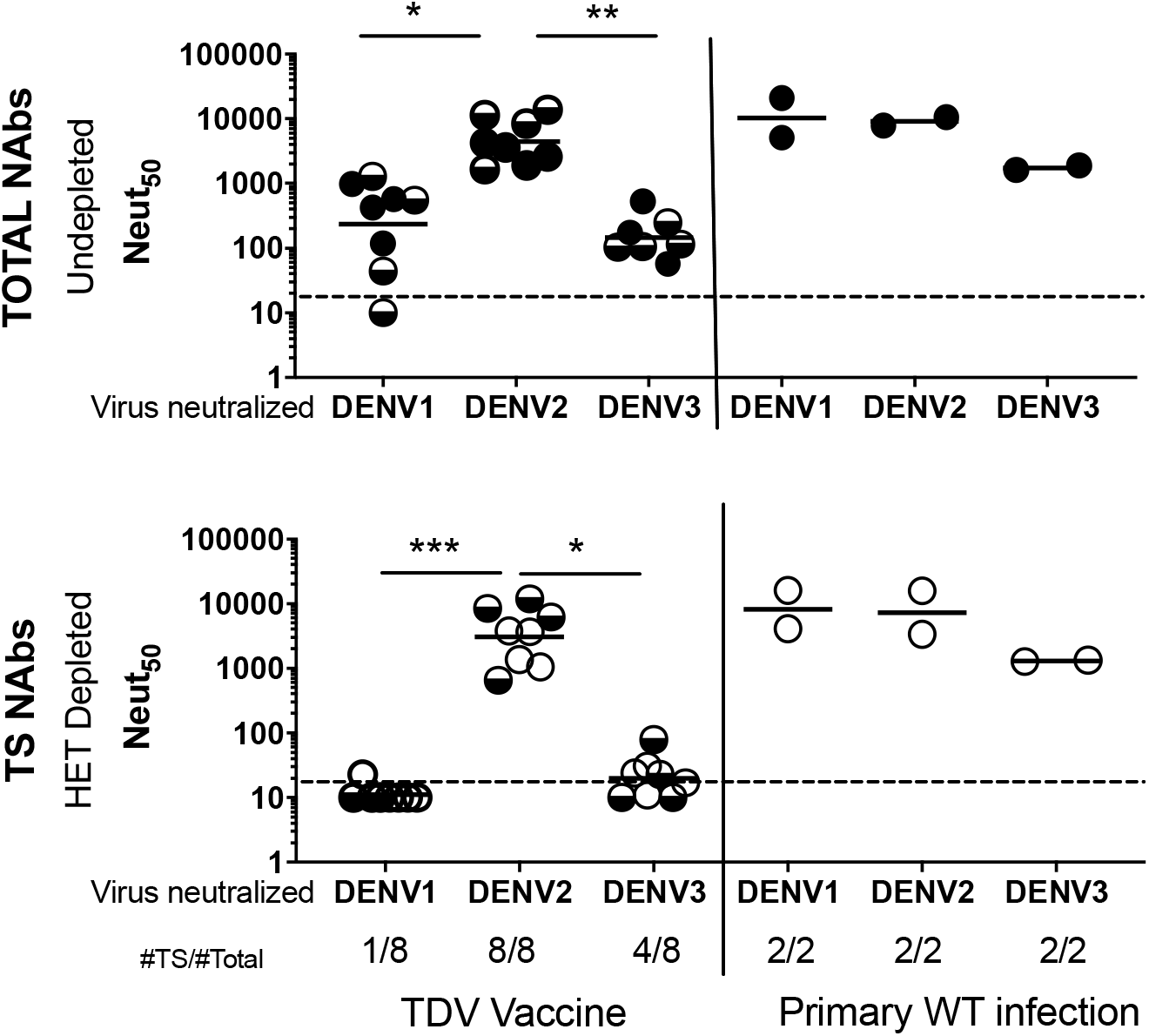
50% Neutralization titers (Neut_50_) in undepleted (A) and Heterologous virus-depleted (B) baseline seronegative NHP vaccinated with TDV (left) or infected with WT DV1, DV2 or DV3 (right). Non-human primates were vaccinated with one dose (filled circles in (A), empty circles in (B) or two doses (half filled circles in (A) and (B) of TDV, or infected once with wild type DV1, DV2 or DV3. Animals were bled 180 days after infection or last vaccination. Each sample was BSA-depleted (Undepleted control) (A) or depleted of cross-reactive NAbs using heterologous serotype viruses (HET depleted) (B). Neut50 titers to DV1, DV2 and DV3 were determined by mFRNT assay. Each dot represents one animal. Line represents the GMT of the Neut_50_ titers. The lowest serum dilution tested was 20. Statistical analysis was done using nonparametric Friedman test with Dunn’s multiple comparison among serotypes after TDV vaccine.

We conclude that NHPs infected with WT DENV1, 2 or 3 develop high levels of NAbs that mainly target type-specific epitopes on each serotype. TAK-003 induced a similar type-specific response to DENV2. The vaccine induced low levels of NAbs to DENV1 and 3 that mainly targeted epitopes that were conserved between serotypes. In this NHP study, the response to DENV4 was not evaluated.

### Levels of DENV serotype specific NAbs in dengue-naïve humans after TAK-003 vaccination

We obtained 30 vaccine immune sera from DENV naïve individuals enrolled in Takeda’s DEN-205 study, who received the vaccine (S2 Table). The samples were collected 180 days after vaccinating adults with one dose of TAK-003. NAbs to DENV1, 2, 3 and 4 were detected in 73% (GMT=57), 93% (GMT=474), 70% (GMT=90) and 57% (GMT=44) of subjects respectively (Fig 2A). The overall NAbs titers to DENV2 were significantly higher than those to the other 3 serotypes. Subjects had received one of two formulations, TDV (N=14) or HD-TDV (N=16), that differed in the dose of the TDV2 component, which was 10 times higher in TDV-HD. Unless otherwise indicated, data from both formulations were grouped together for analysis. When the two formulations, TDV and HD-TDV, were compared, we found that HD-TDV induced significantly higher Neut_50_ titers against DENV2 and higher levels of TS NAbs, while no differences were found for DENV1, DENV3 and DENV4 (S4 Table). Fig 2B shows the NAb titers after HET depletion, providing a measure of TS NAbs stimulated by each vaccine component. Out of 22 DENV1 responders (subjects with DENV1 Neut_50_ titers >20), 15 subjects failed to neutralize DENV1 after cross-reactive Abs were depleted. Out of 28 subjects with DENV2 Neut_50_ titers >20, the majority (23) showed neutralization after depletion of CR Abs. DENV3 and DENV4 neutralization showed a pattern similar to DENV1, where most subjects failed to neutralize the corresponding serotype after removal of cross-reactive antibodies (Fig 2B). As controls, we analyzed immune sera from individuals exposed to past primary DENV1 (6 subjects) or DENV2 (8 subjects) infections before enrolling in study 205. All 14 subjects had high levels of TS NAbs to DENV1 or 2 (Fig 2A, 2B, right panels).

**Figure 2.**
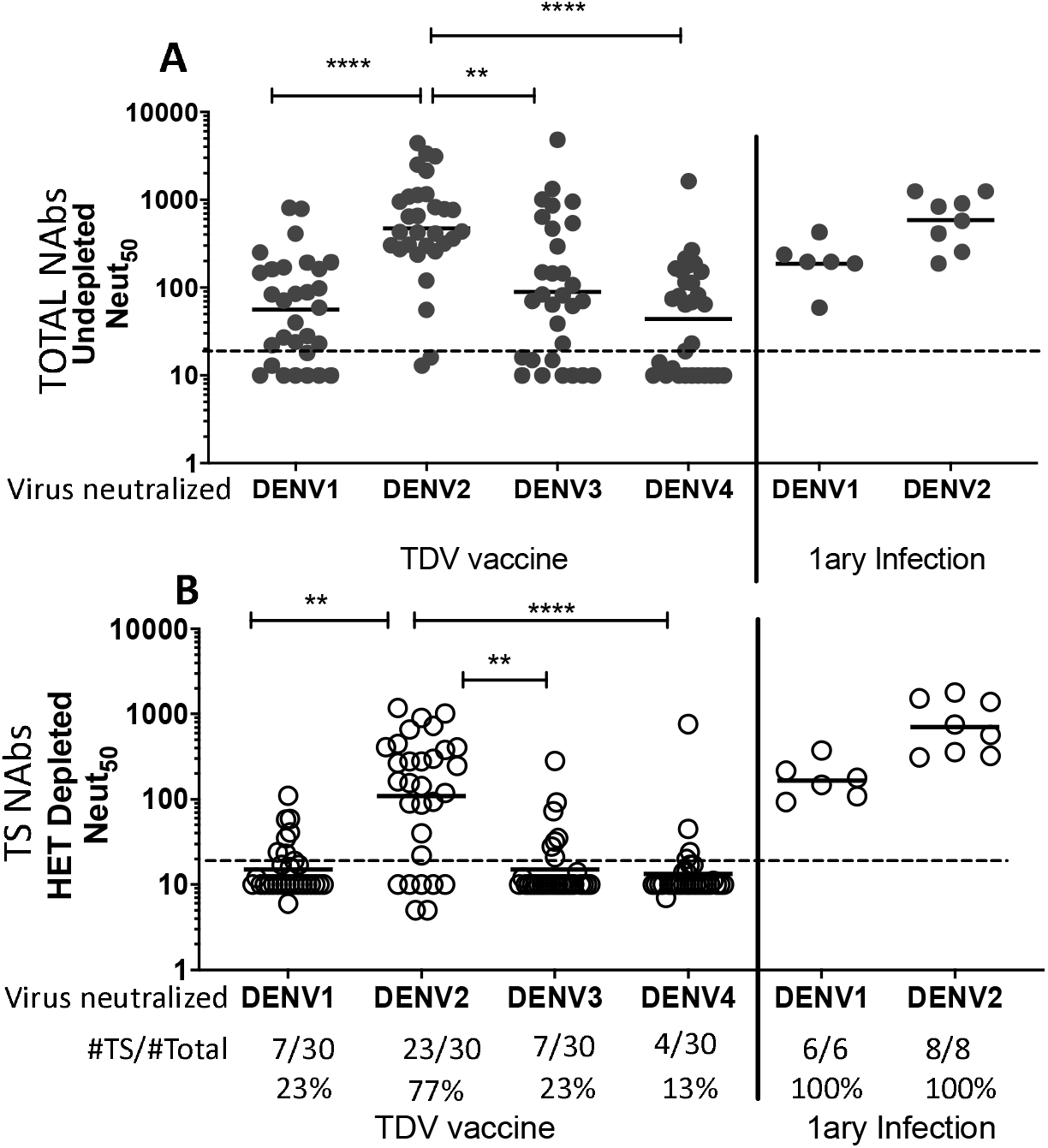
50% Neutralization titers (Neut_50_) in undepleted (A) and Heterologous virus-depleted (B) baseline seronegative subjects vaccinated with one dose of TDV (left) or after exposure to primary DV1 or DV2 infection (right). Sera from 30 subjects were collected 180 days after TDV vaccination. Each sample was BSA-depleted (Undepleted control) (A) or depleted of cross-reactive NAbs using heterologous serotype viruses (HET Depleted) (B). To measure DV1 and DV3 TS neutralization, HET depletion was done with a mix of DV2+DV4 dynabeads. For DV2 and DV4 neuts, Het depletion was done with a mix of DV1+DV3 dynabeads. Each value represents the titer after subtraction of background homologous depletion titer. Each dot represents one subjects. Line represents the GMT of the Neut_50_ titers. The # of subjects with evidence of TS NAbs >20 after HET depletion/#total subjects analyzed is shown at the bottom of panel B. The lowest serum dilution tested was 20. Statistical analysis was done using nonparametric Friedman test for multiple comparison among serotypes after TDV vaccine.

We also performed a more stringent analysis to estimate levels of TS NAbs that took into consideration the efficiency of antibody depletion. In some individuals with high titers of antibodies after vaccination (mostly DENV2 Abs), it was difficult to remove all antigen-specific antibodies using the beads coated with purified virions. As incomplete removal of cross-reactive antibodies targeting epitopes conserved between serotypes can result in overestimating levels of TS NAbs, we only used samples that achieved >90% removal of cross reactive antibodies for calculating TS responses. The number of samples that met the acceptance criterium were 20/30 for DENV1, 29 for DENV2, 26 for DENV3 and 26 for DENV4. Fig 3 shows the Neut_50_ titers after BSA control depletions only for those samples that met the acceptance criteria for TS analysis. We found DENV1 TS NAbs in 1/20 (5%), DENV2 TS NAbs in 24/29 (83%), DENV3 TS NAbs in 3/26 (12%) and DENV4 TS NAbs in 7/27 (27%). In contrast, all the control subjects exposed to WT primary DENV1 or DENV2 infection developed high levels of TS NAbs (Fig 3).

**Figure 3.**
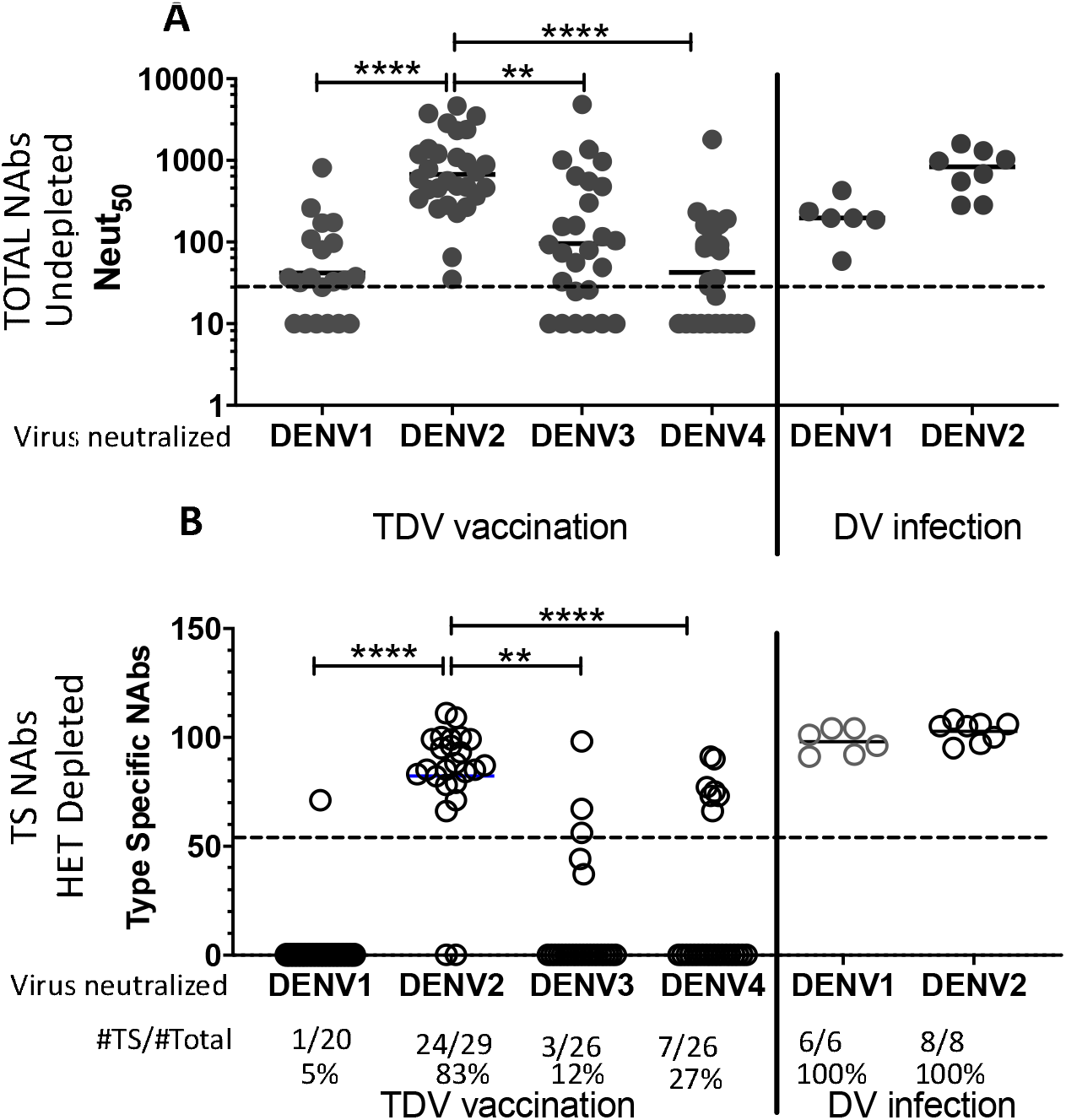
Analysis of the levels of TS Nabs in a subset of DEN-205 clinical samples that met enhanced inclusion criteria for TS analysis. Samples were included in this analysis only if the efficiency of Ab depletion for the serotype used as heterologous was >90%. Only 20 subjects met the criterium for DV1 analysis, 29 for DV2, 26 for DV3 and 26 for DV4. (A) Total NAb titers in undepleted sera. (B) Estimated levels of TS NAbs using the formula [(log10 Neut_50_ heterologous depletion/log10Neut_50_ BSA depletion)x100]. The dotted line represents the set cutoff value (55) for evidence of TS NAbs. The number of samples with TS NAbs for each serotype is indicated below each graph as a ratio over the total # of samples analyzed. Statistical analysis was done using nonparametric Friedman test with Dunn’s multiple comparison among serotypes after TDV vaccine.

In summary, the results generated by both analyses (Fig 2 and 3) are consistent and show that in humans TAK-003 reliably elicited TS NAbs to DENV2 (77-83%), while TS NAbs to DENV1, DENV3, and DENV4 were detected at lower levels in a subset of individuals (5-27%) who were vaccinated.

### Mapping the DV2 TS NAb response in NHP and in humans

Having established that TAK-003 reliably induced TS NAbs to DENV2 in people and monkeys, next we mapped epitopes on DENV2 recognized by those TS NAbs. Previous studies by our group have demonstrated that some TS NAbs elicited after DENV2 natural infection map to quaternary antigenic regions/epitopes involving EDIII of the E protein (29, 33). We used a recombinant chimeric rDENV4/2 virus displaying the DENV2 TS epitopes centered on EDIII on a DENV4 E protein backbone to map vaccine responses. Gain of neutralization by the chimeric DENV4/2 virus over the parental DENV4 backbone was evidence of DENV2 neutralizing Abs targeting sites in EDIII. The properties of the chimeric and parental viruses used in this experiment are described elsewhere (29, 33).

Sera from NHP were tested for neutralization against DENV4/2, DENV2 and DENV4 viruses (Fig 4A). In order to assess gain of neutralization to the transplanted epitope in the chimeric virus, we had to deplete the sera of DENV4 binding Abs to just measure responses directed to the transplanted DENV2 domain. After removal of DENV4 binding Abs, the vaccinated monkeys maintained high titers to DENV2 (Neut_50_ GMT=3077), did not neutralize DENV4 (Neut_50_ < 10) and efficiently neutralized the DENV4/2 virus chimera (Neut_50_ GMT=2223) (Fig 4A). A similar pattern was observed in 2 monkeys infected with WT DENV2 (Fig 4B), indicating both groups developed DENV2 TS Abs that target EDIII. Sera from 15 vaccinated humans were tested for neutralization of DENV2, DENV4 and DENV4/2 viruses (Fig 4C). As most subjects had very low levels of DENV4 NAbs, we did not remove DENV4 binding antibodies before performing the neutralization assays. Twelve of the 15 subjects tested neutralized DENV4/2 (GMT=104) better than DENV4 alone, further implicating EDIII as a target of neutralizing Abs. We concluded that most subjects vaccinated with TAK-003 develop DENV2 TS NAbs that map to epitopes centered on EDIII, which is a major target of NAbs induced by WT DENV2 infections (29, 33).

**Figure 4.**
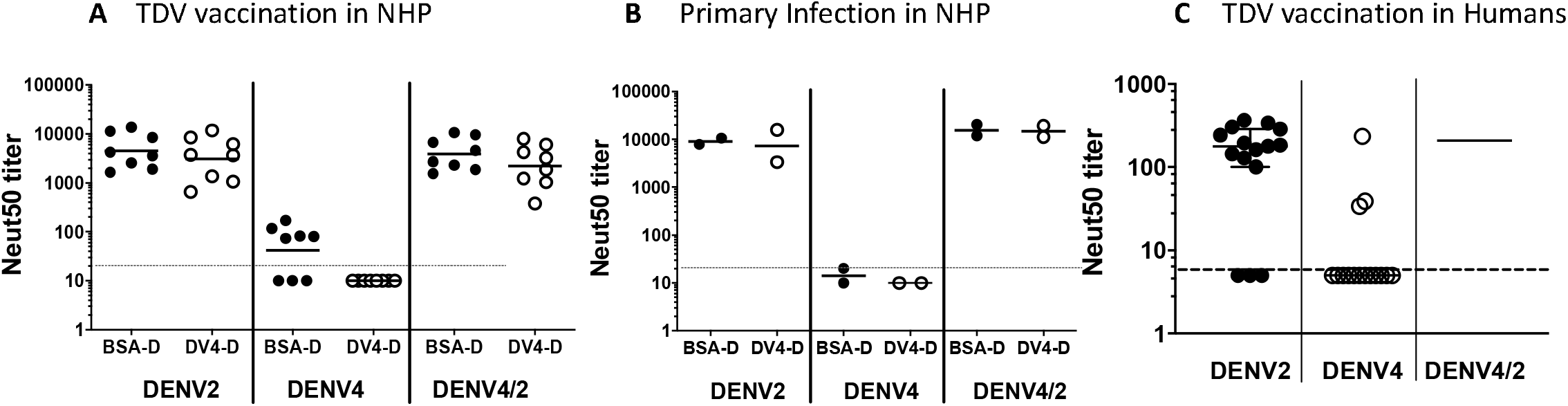
Tracking the molecular specificity of DV2 TS NAbs to epitopes on EDIII in sera from WT DV2 infected NHPs (A), TDV vaccinated NHPs (B) or TDV vaccinated baseline seronegative humans (C). Neut_50_ titers to parental viruses DV2 (EDIII epitope donor), DV4 (backbone recipient of epitope transplant) and chimeric virus DV4/2EDIII (DV2 EDIII transplanted into DV4 E protein) were determined in sera collected 180 days after infection or vaccination. (A) and (B) show titers in undepleted (BSA-D) and DV4-depleted sera. (C) shows titers in undepleted sera. Horizontal lines and error bars represent GMT +/- 95% CI. Dotted line represents lower limit of detection.

## Discussion

The development of tetravalent dengue vaccines has been guided by Nabs to each serotype as a correlate of safe and effective vaccine induced immunity. However, the presence of Nabs alone to each serotype has proven to be an unreliable correlate of protection (18). In baseline dengue seronegative individuals who receive live-attenuated tetravalent DENV vaccines, TS-Nab, which is a measure of immunity independently stimulated by each vaccine component, may be a better correlate of protection.

In a previous study, we analyzed specimens from Takeda studies 104 and 203 to understand the specificity of vaccine induced serum Nabs in individuals who were DENV seronegative or seropositive at baseline (34). In individuals who were seronegative at baseline, we observed high levels of DENV2 Nabs that tracked with TS epitopes on serotype 2. Our results were in agreement with the DENV2 vaccine component replicating and independently stimulating immunity in study subjects. The results for DENV1, 3 and 4 were mixed and harder to interpret. Overall levels of DENV1, 3 and 4 Nabs were about 10-fold lower than the DENV2 titers. Moreover, only about half the study subjects had sufficient levels of DENV1, 3 or 4 Nabs for mapping the responses. Mapping studies with epitope transplant chimeric DENVs and Ab blockade assays indicated that a few individuals did develop Nabs that recognized unique epitopes on DENV1 or 3. However, these conclusions were subject to alternate explanations because we did not use Ab depletion strategies to isolate and characterize TS serum Abs. To overcome this ambiguity about the specificity of TAK-003 induced DENV1, 3 and 4 neutralizing antibodies, in the current study we used antibody depletion methods to remove DENV cross-reactive antibodies before determining the level and specificity of type-specific antibodies.

In the current analysis of 30 seronegative subjects who received TAK-003, most subjects (83%) had high levels of DENV2 TS-Nabs that tracked with known epitopes on DENV2. In contrast, the Nab levels to the other three serotypes were lower and 5%, 12% and 27% of subjects had TS Nabs to DENV1, 3 and 4 respectively. NHPs that received TAK-003 had high levels of DENV2 TS NAbs and low or no detectable TS NAbs to DENV1 or 3. Together, these results indicate that TAK-003 stimulates an independent and type-specific DENV2 NAb response in subjects who were seronegative at baseline. In contrast, most subjects developed lower levels of mainly DENV cross-reactive NAbs to the other three serotypes that are likely derived from the dominant DENV2 vaccine component. While the overall NAb titers to DENV4 were the lowest compared to the other serotypes, when individuals did develop DENV4 Nab titers >40, the response often included TS-Nabs (58%). This observation suggests that low levels of DENV4 Nabs that develop after vaccination may sometimes originate from the DENV4 vaccine component.

Our conclusion that antibodies stimulated by TAK-003 are mainly derived from the dominant DENV2 is supported by other studies to characterize the vaccine in animals and humans (26, 27) (28). Soon after vaccination, investigators have readily detected replicating DENV2 vaccine virus or viral RNA and rarely detected the other three vaccine componets (27) (28). More recently, Biswal et al. analyzed Nab levels in 702 DENV seronegative subjects who received two doses of the vaccine and observed that DENV2 Nab titers were 10 fold or more higher than the DENV1, 3, 4 titers (24).

Phase III clinical trials of TAK-003 are currently ongoing. Results of vaccine efficacy at 12 and 18 months after vaccination have been published (24, 25). At 18 months, overall vaccine efficacy (VE) in individuals seronegative at baseline was 66%. However, VE differed by serotype. There was significant VE to DENV1 and DENV2, lack of efficacy to DENV3 and inconclusive results to DENV4. While more data are needed to understand long term safety and efficacy, current Phase III clinical studies provide the opportunity to study the roles of TS and CR NAbs and T cell immunity in durable immunity.

## Ackowledgements

Ramesh Jadi and Matt de la Cruz contributed in the preparation of purified DENV antigen for depletions.

## Supporting information

**S1 Table. Source of nonhuman primates serum samples**

**S2 Table. Source of human serum samples**

**S3 Table. Takeda TAK-003 vaccine formulations used**

**S4 Table**. Neut_50_ **titers induced after one dose of TDV or HD-TDV in dengue naïve adults**

