## Supplemental Table 1 for "Defining levels of dengue virus serotype-specific neutralizing antibodies induced by a live attenuated tetravalent dengue vaccine (TAK-003)"

### S1 Table. Source of nonhuman primate serum samples

| Study | Serotype Analyzed | Treatment Group | N | Vaccination or infection day | Day of serum collection |
|---|---|---|---|---|---|
| Takeda study DNHP007 Tetravalent TDV vaccine | DV1 | One dose TDV | 4 | 0 | 180 |
| | | Two doses TDV | 4 | 0, 180 | 180 post dose 2 |
| | | WT DV1 infection | 2 | 0 | 180 |
| | DV2 | One dose TDV | 4 | 0 | 180 |
| | | Two doses TDV | 4 | 0, 180 | 180 post dose 2 |
| | | WT DV2 infection | 2 | 0 | 180 |
| | DV3 | One dose TDV | 4 | 0 | 180 |
| | | Two doses TDV | 4 | 0, 180 | 180 post dose 2 |
| | | WT DV3 infection | 2 | 0 | 180 |
