## Supplemental Table 2 for "Defining levels of dengue virus serotype-specific neutralizing antibodies induced by a live attenuated tetravalent dengue vaccine (TAK-003)"

**S2 Table. Source of human serum samples**

| Study | Serostatus at vaccination | Vaccine (One dose) | N | Day of serum collection |
|---|---|---|---|---|
| Takeda study DEN-205 Phase 2 trial Singapore Adults 21-45 Years Old (N=400) | Dengue seronegative | TV TDV | 14 | 180 days post vaccination |
| | | TV HD-TDV | 16 | 180 days post vaccination |
| | Dengue 1 seropositive | - | 7 | Day 0 pre-vaccination |
| | Dengue 2 seropositive | - | 9 | Day 0 pre-vaccination |
