## Supplemental Table 3 for "Defining levels of dengue virus serotype-specific neutralizing antibodies induced by a live attenuated tetravalent dengue vaccine (TAK-003)"

### S3 Table. Takeda Vaccine formulations used

| Formulation | FFU/dose | | | |
|---|---|---|---|---|
| | TDV1 | TDV2 | TDV3 | TDV4 |
| TDV* | $2 \times 10^4$ | $5 \times 10^3$ | $1 \times 10^5$ | $3 \times 10^5$ |
| HD-TDV | $2 \times 10^4$ | $5 \times 10^4$ | $1 \times 10^5$ | $3 \times 10^5$ |

* Used in ongoing Phase 3 trials
