## Supplemental Table 4 for "Defining levels of dengue virus serotype-specific neutralizing antibodies induced by a live attenuated tetravalent dengue vaccine (TAK-003)"

### S4 Table. $Neut_{50}$ titers induced after one dose of TDV or HD-TDV in dengue naïve adults

| | DV1 | | DV2 | | DV3 | | DV4 | |
|---|---|---|---|---|---|---|---|---|
| | TDV | HD-TDV | TDV | HD-TDV | TDV | HD-TDV | TDV | HD-TDV |
| | 69 | 181 | 366 | 2844 | 171 | 94 | 161 | 125 |
| | 98 | 263 | 283 | 459 | 80 | 481 | 79 | 162 |
| | 421 | 205 | 789 | 891 | 955 | 1360 | 193 | 88 |
| | 174 | 172 | 271 | 1187 | 555 | 49 | 234 | 93 |
| | 94 | 802 | 2319 | 4642 | 4857 | 160 | 163 | 192 |
| | 81 | 224 | 467 | 3503 | 117 | 650 | 605 | 329 |
| | 819 | 109 | 2386 | 1203 | 970 | 304 | 1817 | 93 |
| | 38 | 157 | 66 | 1395 | 74 | 155 | 10 | 259 |
| | 28 | 50 | 338 | 729 | 104 | 1013 | 10 | 36 |
| | 10 | 10 | 35 | 952 | 10 | 10 | 10 | 10 |
| | 32 | 10 | 474 | 227 | 72 | 26 | 85 | 10 |
| | 10 | 34 | 483 | 1090 | 25 | 10 | 29 | 22 |
| | 95 | 33 | 41 | 601 | 33 | 80 | 10 | 33 |
| | 10 | 37 | 564 | 3754 | 10 | 56 | 10 | 10 |
| | | 10 | | 257 | | 10 | | 10 |
| | | 37 | | 440 | | 10 | | 10 |
| GMT | **61** | **71** | **336** | **1032** | **123** | **90** | **65** | **47** |
